# Rational Drug Design of Targeted and Enzyme Cleavable Vitamin E Analogs as Neoadjuvant to Chemotherapy: *In Vitro* and *In Vivo* Evaluation on Reduction of Cardiotoxicity of Doxorubicin

**DOI:** 10.1101/2021.05.20.445072

**Authors:** Raghu Pandurangi, Orsolya Cseh, H. Artee Luchman, Siguang Xu, Cynthia Ma, Sanjeewa N. Senadheera, Laird Forrest

## Abstract

Traditional drug design focus on specific target (s) expressed by cancer cells. However, cancer cells outsmart the interventions by activating survival pathways and/or downregulating cell death pathways. As the research in molecular biology of cancer grows exponentially, new methods of drug designs are needed to target multiple pathways/targets which are involved in survival of cancer cells. Vitamin E analogues including a-tocopheryl succinate (α-TOS) is a well-known anti-tumoregenic agent which is well studied both *in vitro* and *in vivo* tumor models. However, lack of targeting cancer cells and unexpected toxicity along with the poor water solubility of α-TOS compelled a rational drug design using both targeting and cleavable technologies incorporated in the new drug design. A plethora of Vitamin E derivatives (AMP-001, 002 and 003) were synthesized, characterized and studied for the improved efficacy and lowered toxicity in various cancer cells *in vitro*. Preliminary studies revealed AAAPT leading candidates reduced the invasive potential of brain tumor stem cells, synergized with different drugs and different treatments. AAAPT leading drug AMP-001 enhanced the therapeutic index of front-line drug Doxorubicin in triple negative breast cancer (TNBC) tumor rat model preserving the ventricular function when used as a neoadjuvant to Doxorubicin. These results may pave the way for reducing the cardiotoxicity of chemotherapy in clinical settings.

## Introduction

Conventional drug designs are based on the identification of specific biological targets involved in the progression of disease, varied target expression in patients, binding of the potential drug to target and modification of the target post binding of the drug^1^. Recent investigations on the molecular biology of cancer open a new type of targets namely, pathways involved in the tumorigenesis^2^. For example, cancer cells enhance the survival pathways and/or enzymes (e.g. NF-kB pathway, PARP) and inhibit or downregulate the cell death pathways (e.g. CD95, ASK1) to desensitize themselves to intervention, irrespective of the nature of intervention^3^. Unfortunately, the same pathways are involved in normal cell growth, despite the levels of enzymes or activating/inhibiting potential of pathways may vary to a great extent. The concept of drug targeting using tumor specific biomarkers is not new for oncology drugs. However, targeting the dysregulated pathways involved in the desensitization process is still relatively new.

α-tocopheryl derivatives are class of compounds which showed interesting biological activities for cancer treatment^4^. Particularly, α-tocopheryl succinate (α-TOS) has shown tremendous anti-tumorigenic potential both *in vitro* and *in vivo* tumor models^5^. However, on extending the promising work of α-TOS to immunocompetent tumor model which could be translational to clinical product lack the required safety profile for commercialization^6^. For example, studies in an immunocompetent mouse *in vivo* model showed that α-TOS was not only ineffective at the published doses but also resulted in severe side effects due to lack of targeting^6^. α-TOS also faced problems with water solubility, bioavailability and formulation issues demanding encapsulation of the drug into liposomes^7^. The lack of proof of targeting, particularly *in vivo* appears to be responsible for the lack of clinical product so far, for α-TOS. Here, we report various derivatives of α-TOS (antioxidant) incorporating targeting vector, cleavable linker and pegylation to improve the pharmacokinetic parameters required for clinical applications. Our studies show an immense improvement in targeting cancer cells by AAAPT drugs, particularly cancer stem cells (CSCs) and cancer resistant cells (CRCs) which are presumably responsible for the treatment failure and recurrence of cancer^8^. The leading drug is also designed with two linkers cleavable by low pH^9^ and Cathepsin B^10^ which is overexpressed in several cancers.

Triple negative breast cancer (TNBC) is an aggressive disease, which disproportionately accounts for nearly half of all breast cancer-related deaths, compared to 15-20% of breast cancer patients^11^. Occurring primarily in women younger than 50 years of age, TNBC prevalence is highest in patients of African American and Hispanic descent^12^. The mortality rate for TNBC patients is 40% within 5 years of diagnosis^13^. Patient survival after cancer recurrence, within 3-5 years of diagnosis rarely extends beyond 12 months^14^. Without effective treatment strategies, roughly 20,000 of TNBC patients die yearly in the U.S. alone, making the development of potent TNBC therapies an urgent medical need^15^.

TNBC is typically classified as grade 3 are metastatic in nature in 66% of patients, with large tumors and metastatic spread to visceral organs, lung or brain already present^16^. TNBC cells lack the expression of biomarkers, such as estrogen receptor (ER), progesterone receptor (PR) or human epidermal growth factor receptor HER-2, which are used in the design of potent targeted therapies. Hence, current targeted therapy may not work efficiently for TNBC patients^17^. Cellular and molecular heterogeneity is another important hallmark of TNBC tumors, further hindering the design of targeted therapies^18^.

Together, these factors warrant current use of cytotoxic front-line chemotherapeutics as primary neoadjuvant, adjuvant and metastatic treatment modality for TNBC^19^. Chemotherapy, particularly anthracyclines are still the first line of treatments for cancer, despite high off-target toxicity, including well reported cardiotoxicity, induction of stemness to expand cancer stem cells (CSCs), eliminate bone marrow cells and may have to be stopped due to lymphedema^20^. This is a serious clinical problem which results in an increase in recurrence rate (13% for kidney cancer, 36% for breast and almost 100% for brain cancer) and make tumors refractory to future treatments^21^. The risk of anthracycline related cardiomyopathy increases with a higher cumulative anthracycline dose^22^. About 3% of the dose persist even after the completion of therapy leading to heart failure and death for a dose 400 mg/m^2^, 7% for a dose of 550 mg/m^2^, and 18% for a dose of 700 mg/m^2^. Chronic anthracycline related cardiotoxicity (ARC) and subsequent adverse sequelae was reported to be around 50 % of all the cases^23^. Reversal of cardiotoxicity is possible with a combination of oncology and cardiology drugs <u>provided an early diagnosis is made</u> (e.g. poly (ADP-ribose) polymerase (PARP) inhibitors^24^, NF-kB inhibitors^25^ and some natural antioxidants^26^). Hence, predicting ARC and reversal of cardiotoxicity using a combined formulation of drugs *in vivo* is a top priority in the management of cancer patients which can lead to quality of life, particularly for triple negative breast cancer (TNBC) patients who have no option, but to rely on chemotherapy, despite high off-target toxicity^27^.

Here we want to report the synthesis and biological evaluation of leading AAAPT dugs in vitro in various types of cancer cells and to show that these derivatives are synergistic with the current treatments (e.g. doxorubicin, paclitaxel), particularly for TNBC treatment. AMP-001 showed a significant potential to reduce the invasive potential of brain tumor stem cells (BTSCs). We have also investigated the anti-tumorigenic potential of AMP-001 *in vivo* using rat TNBC model where the synergistic potential of AMP-001 with doxorubicin resulted in lowering the Doxorubicin dose by 50 % yet inducing significant tumor regression and preserving ventricular function compared to Doxorubicin alone.

## Material and Methods

Synthesis of the four leading drugs will be described. All precursor chemicals were bought from Sigma and used as such using MSDS data.

### Synthesis of AMP-001. AMP-002 and AMP-003

AAAPT drugs have three parts: a) α-tocopherol part, alkylation part and dipeptide valine-citrulline (Scheme 1). The precursor of all AMP class of compounds is α-tocopherol which was modified by an alkyl chain either with ether link (AMP-002) or with ester link (AMP-003). AMP-001 was pegylated after alkylating OH group of tocopherol before conjugating with valine-citrulline di peptide linker.

**Scheme 1.**
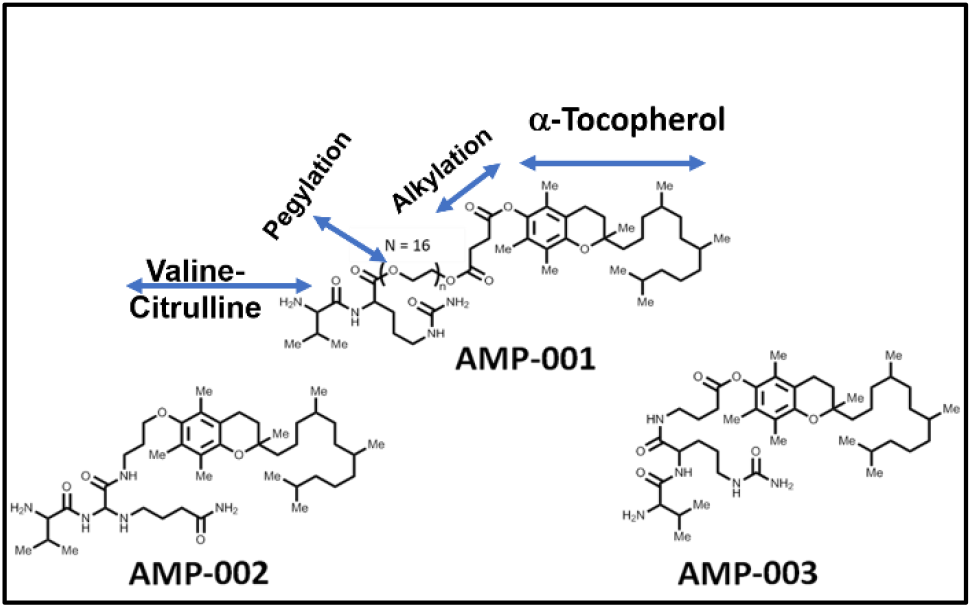

#### A) Alkylation of α-tocopherol with Ether link to get Tocopheryl oxy propyl amine (TOPA)

##### Step 1: Tert-butyl (3-bromopropyl)carbamate 2

To a stirred suspension of **compound 1** (2.5 g, 11.46 mmol) in Dry DCM (25 mL) was added TEA (5.6 mL, 40.11 mmol) at 0° C under inert atmosphere. To this Boc anhydride (2.5 mL, 11.46 mmol) was added in drop wise over the period of 15 min. The reaction mixture was stirred at RT for 6 h. After completion of the reaction (by TLC), the reaction mixture was diluted with ice-cold water (75 mL) and extracted with DCM (3 x 75 mL). The combined organic extracts were washed with water (75 mL), brine (50 mL), dried over anhydrous Na_2_SO_4_ and concentrated under reduced pressure. The obtained crude material was purified by silica gel column chromatography **(SiO2, 60-120 mesh)** (eluent: 10% EtOAc/Hexane) to afford **compound 2** (2.6 g, 10.92 mmol, 95%) as sticky syrup. ^**1**^**H NMR (400 MHz, CDCl_3_):** δ 4.92 (bs, 1H), 3.52 (t, *J* = 7.2 Hz, 1H), 3.27 (t, *J* = 7.5 Hz, 2H), 2.19 (q, *J* = 8.2 Hz, 2H), 1.91(s, 9H). **MS (ESI):** *m/z* 239 [M^+^+1].

**Scheme 2.**
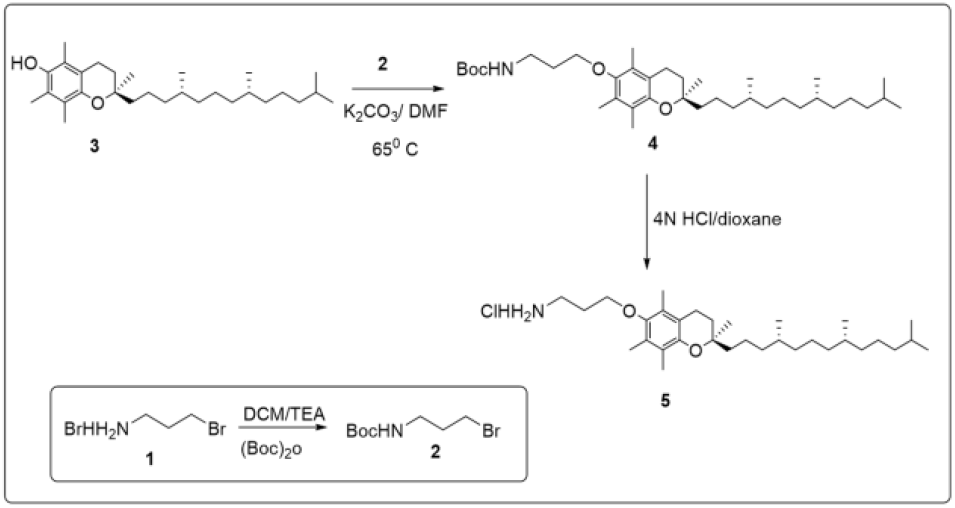

##### Step II. tert-butyl(3-(((R)-2,5,7,8-tetramethyl-2-((4R,8R)-4,8,12-trimethyltridecyl)chroman-6-yl)oxy)propyl)carbamate 4

To a stirred solution of Tocopherol 3 (2.5 g, 5.81 mmol) in dry DMF (15 mL) was added compound 2 (2.49 g, 10.46 mmol) followed by K_2_CO_3_ (2.3 g, 17.43 mmol) at RT under inert atmosphere. The resulting reaction mixture was gradually heated up to 65 °C and stirred for 16 h; progress of the reaction was monitored by TLC. The reaction mixture was diluted with ice-cold water (50 mL) and extracted with EtOAc (3 x 75 mL). The combined organic extracts were washed with water (75 mL), brine (50 mL), dried over anhydrous Na_2_SO_4_ and concentrated under reduced pressure. The crude material was purified by silica gel column chromatography (SiO2, 60-120 mesh) (eluent: 15% EtOAc/Hexane) to afford compound 4 (2.4 g, 4.08 mmol, 62%) as off white solid.

^**1**^**H NMR (5 00 MHz, CDCl3):** δ 3.75 (t, *J* = 8.2 Hz, 2H)), 3.43 (t, *J* = 8.2 Hz, 2H), 2.67 (t, *J* = 7.2 Hz, 3H), 2.35(t, *J* = 7.2 Hz, 2H), 2.19-2.00 (m, 9H), 1.79-1.67 (m, 2H), 1.60-1.48 (m, 3H), 1.43-1.11 (m, 23H), 1.49 (s, 9 H), 0.92-0.82 (m, 12H). **MS (ESI):** *m /z* 588 [M+1]^+^

##### Step 3: 3-(((R)-2,5,7,8-tetramethyl-2-((4R,8R)-4,8,12-trimethyltridecyl)chroman-6-yl)oxy)propan-1-amine hydrochloride 5

To a stirred solution of **compound-4** (2.3 g, 3.91 mmol) in dry 1,4-dioxane (5 mL) was added 4 N 1,4-dioxane solution in HCl (2.3 mL) at 0° C under inert atmosphere. The resulting reaction mixture was stirred at RT for 9 h. After completion of the reaction (by TLC), The resulting mixture was concentrated under reduced pressure to get the sticky syrup, after washing with Ether (HPLC, 2 x 20 mL) to afford fine brown solid as **compound 5** (1.2 g, 2.45mmol, 62%)

^**1**^**H NMR (Varian,400 MHz, CDCl3):** δ 8.35 (bs, 3H), δ 3.35 (τ, *J* = 8.2 Hz, 2H)), 3.43 (t, *J* = 8.2 Hz, 2H), 2.52 (t, *J* = 7.2 Hz, 3H), 2.35(t, *J* = 7.2 Hz, 2H), 2.19-2.00 (m, 9H), 1.79-1.67 (m, 2H), 1.60-1.48 (m, 3H), 1.41-1.10 (m, 23H), 0.90-0.80 (m, 12H). **MS (ESI):** *m /z* 488 [M+1]^+^, **HPLC: 92.99%**.

#### B) Alkylation of α-tocopherol with ester link to get Tocopheryl propyl ester amine (TPEA

##### Step 1: 4-((tert-butoxycarbonyl) amino)butanoic acid 2

To a stirred suspension of 4-aminobutanoic acid 1 (1.5 g, 14.56 mmol) in 1, 4-dioxane (15 mL) was added 1N NaOH solution (1.5 mL) at 0° C under inert atmosphere. To this Boc anhydride (3.1 mL, 14.56 mmol) was added in drop wise over the period of 10 min . The reaction mixture was stirred at RT for 4 h. After completion of the reaction (by TLC), the reaction mixture pH was adjusted to 2, by the addition of aqueous K_2_HSO_4_ solution (10 mL), quenched with ice-cold water (50 mL) and extracted with EtOAc (2 x 75 mL). The combined organic extracts were washed with water (50 mL), brine (50 mL). The organic layer was dried over anhydrous Na_2_SO_4_ and concentrated under reduced pressure to afford compound 2 (2.9 g, 14.28 mmol, 98%) as white sold. ^**1**^**H NMR (500 MHz, CDCl_3_):** δ 4.65 (bs, 1H), 3.25 (d, *J* = 8.2 Hz, 1H), 2.44-2.38 (m, 2H), 1.86-1.78 (m, 2H), 1.41(s, 9H). **MS (ESI):** *m/z* 204 [M^+^+1]

**Scheme 3.**
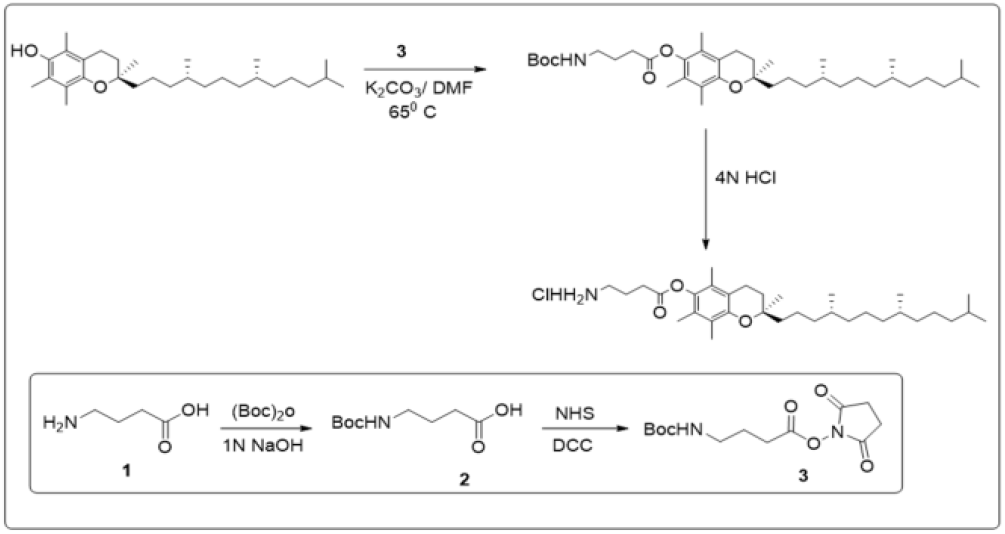

##### Step 2: 2,5-dioxopyrrolidin-1-yl 4-((tert-butoxycarbonyl)amino)butanoate 3

DCC (2.84 g, 13.79 mmol) was added to a solution of compound 2 (2.8 g, 13.79 mmol) and N-hydroxysuccinamide (1.58 g, 13.79 mmol) in 1,4-dioxane (28 mL), and the mixture was stirred at RT for 3 h, under inert atmosphere. After completion of the reaction (by TLC), diluted with 1, 4-dioxane (30 mL) solution, filtered through a pad of celite, obtain filtrate was concentrated under reduced pressure to get the off white solid as compound 3 (3.6 g crude), which was directly used next step with out further purification.

^**1**^**H NMR (500 MHz, DMSO-*d*_*6*_):** δ 3.10 (q, *J* = 8.2 Hz, 1H 2H), 2.69 (t, *J* = 8.2 Hz, 2H), 1.70-1.68 (m, 2H), 1.39(s, 9H). **MS (ESI):** *m/z* 304 [M^+^+1]

##### Step 3: (2S)-2,5,7,8-tetramethyl-2-(4,8,12-trimethyltridecyl)chroman-6-yl4-((tert-butoxycarbonyl)amino)butanoate 5

To a stirred solution of **compound 4** (2.5 g, 5.81 mmol) in dry DMF (15 mL) was added **compound 3** (3.1 g, Crude) followed by K_2_CO_3_ (2.76 g, 20.33 mmol) at RT under inert atmosphere. The resulting reaction mixture was gradually heated up to 65 °C and stirred for 16 h. After completion of the reaction (by TLC), diluted with ice-cold water (50 mL) and extracted with EtOAc (3 x 75 mL). The combined organic extracts were washed with water (75 mL), brine (50 mL), dried over anhydrous Na_2_SO_4_ and concentrated under reduced pressure. The crude material was purified by silica gel column chromatography (SiO2, 60-120 mesh) (eluent: 20% EtOAc/Hexane) to afford **compound 5** (2.2 g, 3.57 mmol, 62%) as off white solid.

^**1**^**H NMR (500 MHz, CDCl_3_)**: δ 4.75 (bs, 1H), 3.30 (s, 2H), 2.66 (t, *J* = 8.2 Hz, 2H), 2.60 (t, *J* = 8.2 Hz, 2H), 2.83 (s, 3H), 2.00 (s, 6H), 1.86-1.72 (m, 3H), 1.59-1.20 (m, 34H), 0.90-0.82 (m, 12H). **MS (ESI):** *m /z* 616 [M+1]^+^

##### Step 4: (2S)-2, 5, 7, 8-tetramethyl-2-(4, 8, 12-trimethyltridecyl) chroman-6-yl 4-aminobutanoate hydrochloride 6

To a stirred solution of compound-5 (2.1 g, 3.41 mmol) in dry 1,4-dioxane (5 mL) was added 4 N 1,4-dioxane solution in HCl (1.5 mL) at 0° C under inert atmosphere. The resulting reaction mixture was stirred at RT for 9 h. After completion of the reaction (by TLC), The resulting mixture was concentrated under reduced pressure, to get the sticky syrup, after washing with Ether (HPLC) to get the fine brown solid as final compound 6 (1.3 g, 2.52 mmol, 74%). ^**1**^**H NMR (Varian,400 MHz, CDCl3):** δ 8.10 (bs, 3H), 2.89 (t, *J* = 7.2 Hz, 2H),), 2.79 (t, *J* = 10 Hz, 2H), 2.57 (t, *J* = 8.0 Hz, 2H), 2.01 (s, 3H), 1.91-1.89 (m, 9H), 1.75 (q, *J* = 7.2 Hz, 2H), 1.40-1.33 (m, 3H), 1.29-1.02 (m, 23H), 0.84-0.80 (m, 12H). **MS (ESI):** *m /z* 516 [M+1]^+^ **HPLC: 91.68%**.

#### C: Conjugation of Alkyl Tocopheryl Derivetives with Dipeptide Valine-Cittrulin to get AMP-001, AMP-002 and AMP-003 respectively

The common intermediate for the preparation of the above three derivatives is Boc-Val-Cit-OH, which has been prepared by the coupling reaction between Boc-Val-NHS and L-citrulline in DME-THF-water in the presence of sodium bicarbonate at room temperature in 50% yield after trituration with isopropyl ether.

##### Step 1: Synthesis of Boc-Val-Cit-OH

Boc-Val-NHS (10 g, 31.8 mmol) in 80 mL of DME was added to a solution of L-Citrulline (5.85 g, 33.4 mmol) in 20 mL of THF and NaHCO3 (2.8 g, 33.4 mmol) in 80 mL of water. The mixture was stirred at room temperature overnight. Aqueous 15 % citric acid (200 mL) was added and the mixture was extracted with10% isopropyl alcohol/ ethyl acetate (2 × 200 mL). The organic extract was washed with water (2 × 200 mL) and the solvent was evaporated under vacuum. The resulting solid was triturated with 200 mL of isopropyl ether to afford the desired product (6.0 g, 50%, Scheme 4). 1H NMR: (CD3OD) δ 0.92 (3H, d, J= 6.7 Hz), 0.97 (3H, d, J= 6.7 Hz), 1.44 (9H, s), 1.51-206 (5H, m), 3.12 (2H, t, J= 6.7 Hz), 3.90 (1H, d, J= 7.0 Hz) and 4.37-4.41 (1H, m).

**Scheme 4.**
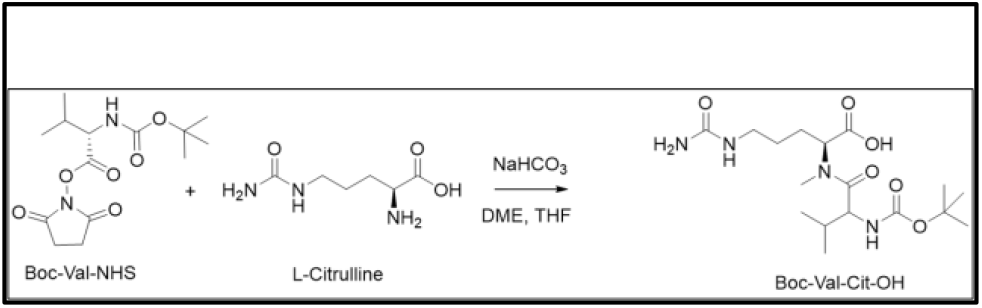

##### II Step: Synthesis of Val-Cit-TPGS (8)

D-α-Tocopherol polyethylene glycol 1000 succinate (TPGS **7**, 506 mg, 0.3 mmol), Boc-Val-Cit-OH **2** (374 mg, 1 mmol), DCC (208 mg, 1 mmol) and DMAP (61 mg, 0.5 mmol) were dissolved in THF (20 mL) and stirred at room temperature for 16 h. The precipitated solid (DCU) was removed by filtration and the filtrate evaporated on rotary evaporator. The residue was purified by silica-gel column chromatography using 10-20 % methanol in dichloromethane as an eluent. Got desired product, **8** (155 mg) as off - white solid.

**Scheme 5.**
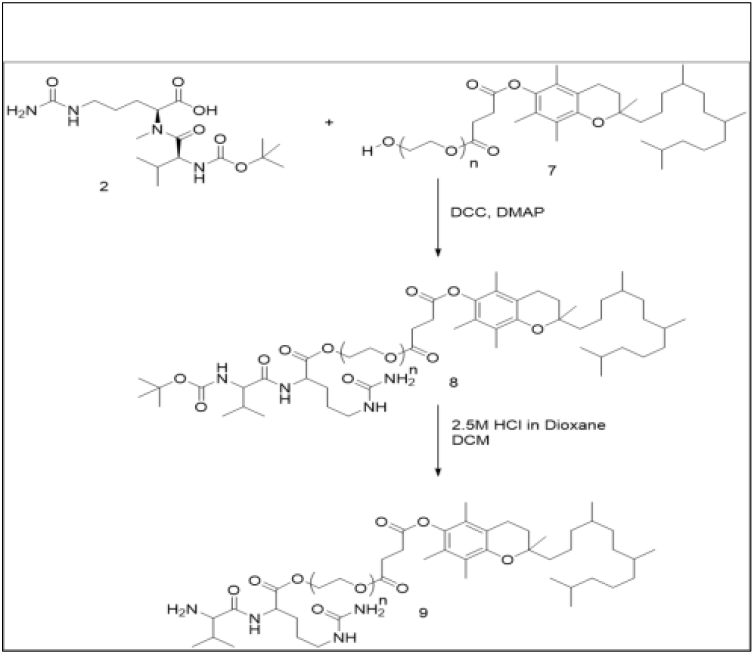

##### Step III: Synthesis of AMP-001 (VC-TPGS 9)

Compound **8** (150 mg) was dissolved in dry DCM (10 mL) and cooled to 0° C. To the resulting solution was added 2.5M HCl in dioxane (3 mL) and allowed to stir at room temperature for 12 h. Then it was concentrated and dried in vacuum oven to get desire product as light brown color solid (103 mg). ^1^HNMR (CDCl_3_) is quite complex. ESI-MS ~ 1942 (Scheme **5**).

#### D: Synthesis of AMP-002

AMP-002 has been synthesized as follows. The reaction of Boc-Val-Cit-OH with TOPA in the presence of EDC and HOBt (1-hydroxybenzotriazole) in THF, gave Boc protected intermediate **2**. Boc group of **2** was removed by treating it with 2.5M HCl in dioxane to afford the desired compound **3**.

**Scheme 6.**
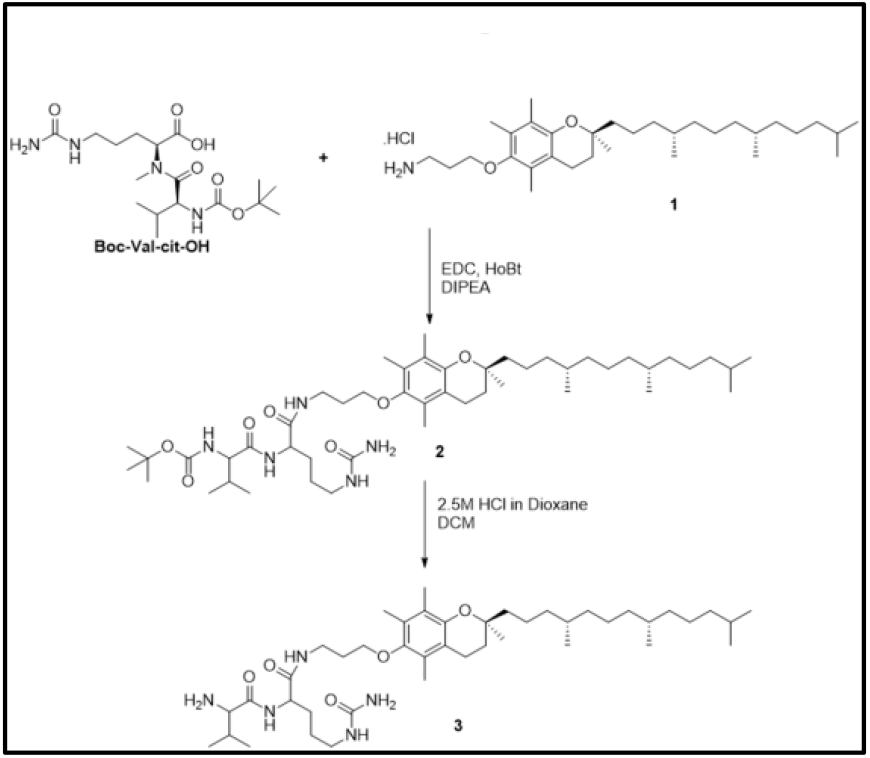

##### Synthesis of AMP-002 (VC-TOPA)

HCl salt of TOPA (**1**, 330 mg, 0.63 mmol), Boc-Val-Cit-OH (330 mg, 0.90 mmol), EDC (200 mg, 1 mmol), HOBt (100 mg, 0.74 mmol) and N,N-diisopropylethylamine (0.3 mL) were dissolved in THF (10 mL) under inert atmosphere. The resulting mixture was stirred at room temperature for 16 h. It was quenched with brine (5 mL) and extracted with DCM (2 × 30 mL). The organic layer separated, dried over Na_2_SO_4,_ concentrated and purified by silica-gel column chromatography using 3-5 % methanol in dichloromethane as an eluent. Got desired product, **2** (200 mg, 37%) as brown color solid. ESI-MS: 844.7 (M+H), Scheme **6**.

#### E: Synthesis of AMP-003: VC-TPEA

##### Step 1: Boc-VC-TPEA

HCl salt of TPEA (**4**, 263 mg, 0.475 mmol), Boc-Val-Cit-OH (263 mg, 0.70 mmol), EDC (193 mg, 1 mmol), HOBt (100 mg) and N,N-diisopropylethylamine (0.3 mL) were dissolved in THF (20 mL) and stirred at room temperature for 16 h. To the resulting mixture was added brine (5 mL) and extracted with DCM (2 X30 mL). The organic layer was dried over Na_2_SO_4,_ filtered, concentrated and purified by silica-gel column chromatography using 3-5 % methanol in dichloromethane as an eluent. Got desired product, **5** (180 mg, 43%) as brown color solid. ESI-MS; 872.6 (M+H).

###### AMP-003: VC-TPEA

Compound **5** (180 mg, 0.20 mmol) was dissolved in dry DCM (10 mL) and cooled to 0°C. To the resulting solution was added 2.5M HCl in dioxane (3 mL) and allowed to stir at room temperature for 12 h. It was concentrated and dried in vacuum oven to get desire product as light brown color hydrochloride salt (150 mg, 94%). %). ^1^HNMR (CDCl_3_): δ 0.96-0.98 (m, 16H), 1.15-1.22 (m, 32H), 1.24-1.44 (m, 4H), 1.51-1.53 (m, 4H), 1.94-2.07 (m, 15H), 2.55 (m, 5H), ESI-MS; 772.6 (M+H).

#### F: Synthesis of Doxorubicin-Tocopherol Conjugate

##### (i) Synthesis of Succinic Acid –Mono-Boc Hydrazine

The structure of the final conjugated is shown in Scheme 2. Succinic anhydride (1 eq.) and DMAP (0.08 eq.) were dissolved in dichloromethane (40 mL). After that, tert-butyl carbamate (1.3 eq.) was added at rt dropwise over 30 min to the reaction mixture under vigorous stirring. After addition was complete, the reaction mixture was stirred overnight at rt. Then, the reaction mixture was concentrated and re-dissolved in ethyl acetate (30 mL). The ethyl acetate layer was washed (20 mL X 3) with dilute HCl solution (pH ~5-6) followed by brine solution (20 mL X 3). Next, ethyl acetate layer was separated and dried over anhydrous MgSO_4_ and concentrated under reduced pressure. The colorless oily product was used in the next step without further purification.

**Scheme 7.**
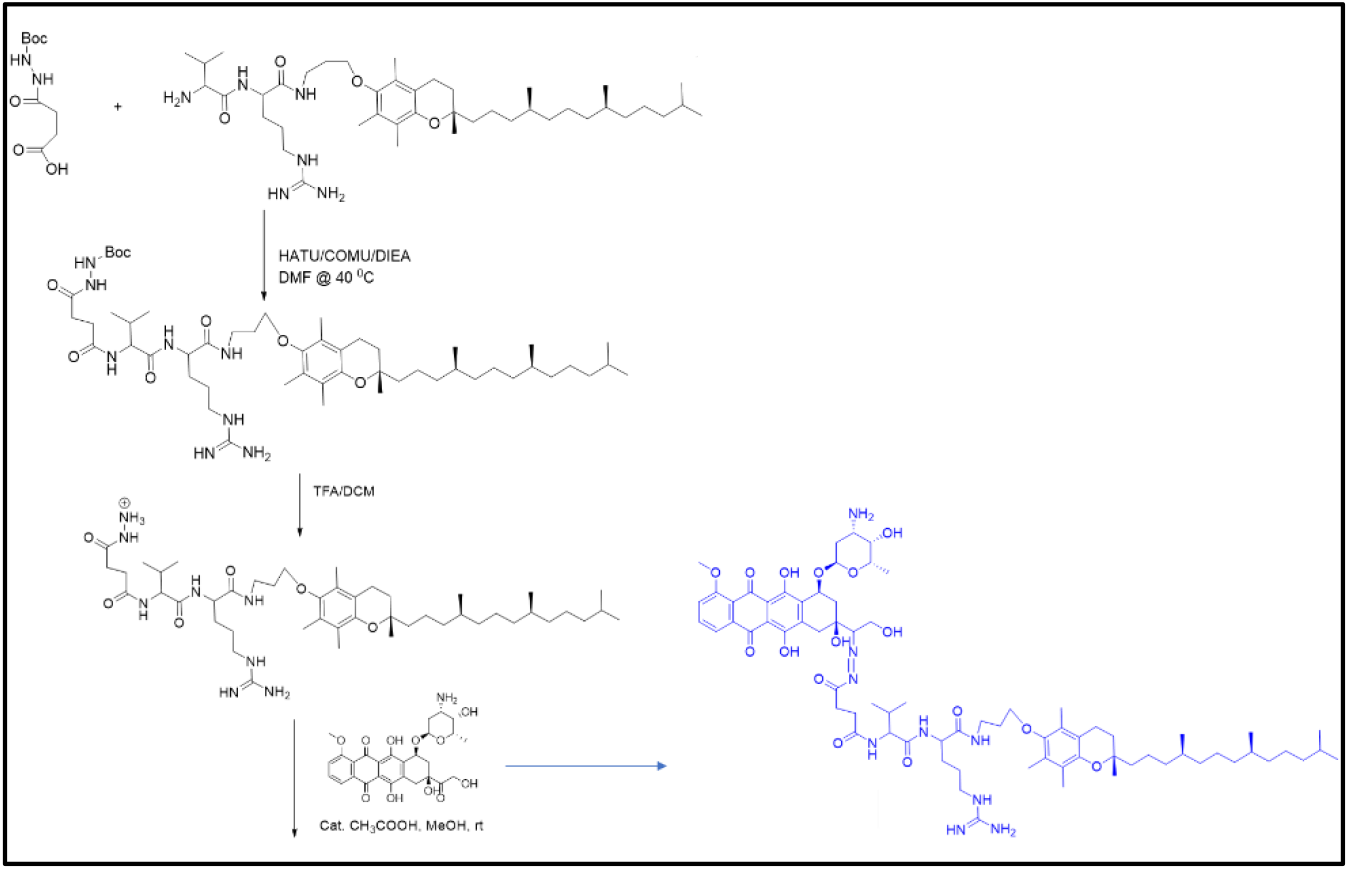

##### (ii) Synthesis of Succinic Acid –Mono-Boc Hydrazine-VA-propyl-Tocopherol conjugate

Succinic Acid –Mono Boc Hydrazine (1.3 eq.) was dissolved in DMF (1 mL) in a scintillation vial. Then, HATU (4 eq.), COMU (4 eq.), and DIEA (10 eq.) were added to the reaction mixture and stirred for 5 min. After that, H-VA-propyl-Tocopherol conjugate (1 eq.) in DMF (1 mL) was added to the reaction mixture dropwise at rt. After addition was complete, reaction mixture was stirred at 40 °C for 24 h. Reaction mixture was then concentrated under reduced pressure. The crude product was triturated with NaHCO_3_ (2 mL x 3) followed by ice cold dilute HCl solution (2 mL x 3; pH ~5-6) and ice cold water (2 mL x 3). The crude product was dried under reduced pressure and was utilized in the next step without further purification.

##### (iii) Synthesis of Succinic Acid –Hydrazine-VA-propyl-Tocopherol conjugate

Succinic Acid –Mono-Boc Hydrazine-VA-propyl-Tocopherol conjugate was dissolved in TFA/DCM (1:1, 2 mL) and was stirred for 4 h. The reaction mixture was concentrated under reduced pressure. The crude was dissolved in purified water (5 mL) was washed with ethyl acetate (5 mL) in a separatory funnel. Water layer was separated, frozen, and lyophilized to obtain the pure product.

##### (iv) Synthesis of Doxorubicin-succinic Acid –Hydrazine-VA-propyl-Tocopherol conjugate

Doxorubicin (1 eq.) and succinic acid –hydrazine-VA-propyl-tocopherol conjugate (1 eq.) and catalytic amount of acetic acid were dissolved in MeOH (2 mL) and stirred at rt for overnight in the dark. Then, the reaction mixture was concentrated under reduced pressure. After that, the doxorubicin-succinic Acid –Hydrazine-VA-propyl-Tocopherol conjugate was precipitated in MeCN/MeOH (5:1) in the dark at 0 °C (the maximum recovery was obtained in the refrigerator at 4 °C overnight). The product was collected by centrifugation (4000 rpm) over 10 min. The supernatant was decanted. The precipitate was washed (x 5) with ice cold solution of MeCN/MeOH (10:1) by centrifugation (finally the supernatant would become a clear solution). Finally, the conjugate was vacuum dried in the dark and analyzed by MS [ESI(+Ve)]; m/z = 468.13 when z = 3 (sodiated).

###### Cell Culture

MDA-MB-231 cells (ATCC HTB-26) were cultured in RPMI 1640 culture medium with L-glutamine (Thermo-Fisher Scientific, Waltham MA, Catalog #11875-093), supplemented with 10% fetal bovine serum and 1% Penicillin/Streptomycin (P/S) solution. Cells were incubated in a humidified incubator with 5% CO_2_. MCF10A (CRL-10317) cells were cultured in DMEM /F12 Ham’s Mixture supplemented with 5% Equine Serum (Thermofisher Catalog # 16050130), EGF 20 ng/ml (Sigma), insulin 10μg/ml (Sigma), hydrocortisone 0.5 mg/ml (Sigma), cholera toxin 100 ng/ml (Sigma), 100 units/ml penicillin and 100 μg/ml streptomycin.

###### Cell Viability Assays

The Cell-Titer-Glo Luminescent Cell viability assay is a method to determine the number of viable cells in culture based on quantitation of the ATP present, as an indicator of metabolically active cells. Cells were seeded at a density of 400 cells / well in 384-well white-walled Cultur-Plate-384 cell culture plates (Perkin-Elmer, Waltham MA, Catalog #6007680), and exposed to test agent the following day. Cells were allowed to grow in the presence of test agent for 72 hours, at which point Cell-Titer-Glo (Promega Corp., Madison WI, Catalog #G7571) reagent was added at a volume equal to the cell culture medium in the plate, according to manufacturer’s instructions. Luminescence was read on an Envision 2104 multilabel reader (Perkin-Elmer, Waltham WI). Cell viability curves were done in quadruplicate for each concentration.

###### Cell Culture on Brain Tumor

All samples were obtained from GBM surgical procedures following informed consent from patients. All experimental procedures were performed in accordance the Health Research Ethics Board of Alberta, Cancer Committee. Brain tumour stem cells (BTSCs) were cultured in neurosphere conditions on non-adherent plates in serum free media supplemented with EGF (20 ng/mL; Peprotech), FGF2 (20 ng/mL; R&D Systems Inc) and heparin sulfate (2 μg/mL; R&D Systems). BTSC lines were confirmed to match their parental primary GBM tumor tissue by short tandem repeat profiling (Calgary Laboratory Services and Department of Pathology and Laboratory Medicine, University of Calgary). Authentication and testing of all cell lines was performed as per American Association for Cancer Research recommendations.

###### Viability assays

For viability assays BTSCs (Bt48, BT67, BT89, BT189) were cultured as neurospheres. To measure the drug effect, dissociated cells were plated in 96-well plates and cells were left untreated, treated with dimethyl sulfoxide (DMSO), AMP100, AMP002, AMP001 or AMP001A in ½ log dose curves ranging from 0.003 to 10 μM re-suspended in DMSO. Following 48 hours or 10-14 days of treatment, cell viability was measured. All cell viability measurements were performed using the alamarBlue™ reagent (Thermofisher) according to the manufacturer’s protocol. All experiments were performed at 3 technical replicates/dose.

###### Invasion Assays

Invasion assays were performed as previously described (Restall et al, J Vis Exp. 2018 Aug 29;(138)). In brief, Small BTSC spheres were collected form culture flasks and treated in 6-well plates. Spheres were treated with DMSO, 0.5 μM AMP001 or 0.25 μM Stattic for 24 hours. Spheres were then transferred to 1.5 mL Eppendorf tubes to allow gravity pellets to form. Media was removed and spheres were resuspended in a Rat Collagen I (Cultrex) matrix. 100 μL/well of collagen suspended spheres was then transferred to a cold 96-well plate with 3 experimental replicates for each treatment condition. Plates were transferred to an Incucyte Live Imaging System (Essen Bioscience). Area of cellular invasion was recorded hourly for 24 hours.

###### Rats Breeding

Breeding pairs were set with one homozygous male (rnu/rnu) and two homozygous females in a cage. Pregnant dams were separated right before new pups were born. Pups are weaned from mothers at 4 weeks of age. Since some breeding nude rats don’t make milk sufficiently on the first pregnancy, pups of first litters rarely survive. Experienced moms produce pups the next pregnancy. For each litter produced after the first litter, 5-8 pups survive to weaning and females usually make up 50% of litter.

###### Cell implantation and tumor measurement

The human breast cancer cell line MDA-MB-231 (ATCC) were grown in DMEM medium, supplemented with 10% (v/v) FBS, penicillin (10 U/mL)–streptomycin (10 U/mL) at 37 °C in humidified 5% CO2 atmosphere. On the day of the cell implantation, MDA-MB 231 cells were trypsinized, washed with PBS and counted per μL. Cells were placed in a sterile microcentrifuge tubes and keep them on ice. Cells were mixed 1:1 with Matrigel (right before the injection) and 12 million cells were injected in the next to last inguinal mammary fat pad in a volume of 100uL. Tumor development was followed (length, width, and height, approximately 1 week after the implantation), Tumor volumes were calculated by the following formula: (1/2 x L x W x H), in which L is the length, W is the width, and H is the height. All rats were housed under a 12-hour light-dark cycle with free access to food and water. This study was performed in accordance with the “Guide for the Care and Use of Laboratory Animals” (2011) of the NIH. The protocol was approved by the Animal Care and Use Committee of the Johns Hopkins Medical Institutions (Baltimore, MD; Animal Welfare Assurance # A-3273-01).

###### DOX and AMP treatment Plan

Rat with comparable sized tumors were randomly divided between the four treatment groups: DOX 5mg/kg, DOX 2.5 mg/kg and AMP-001 75/mg/kg combined, AMP-001 75/mg/kg alone, and no treatment. Doxorubicin was administered via jugular vein injection every 2 weeks for a total of 3 injections. AMP treatment was given intraperitoneally 24 hours before doxorubicin was given.

###### Echocardiography

Transthoracic echocardiography was performed on conscious rat using Vevo 2100 ultrasound scanner equipped with the 250 rat transducer. The rat was introduced into a DecapiCones disposable plastic restraint, plastic on top on the chest removed and pre-warm echo transmission gel was applied to the chest for the imaging. The heart was imaged in a two-dimensional mode followed by M-mode using the parasternal short axis view at a sweep speed of 200mm/sec. After imaging, the ultrasound gel was wiped off and each rat thoroughly dried with disposable towels prior to home cage return. Measurements were acquired using the leading-edge method, according to the American Echocardiography Society guidelines. Left ventricle wall thickness and left ventricle chamber dimensions were acquired during the end diastolic and end systolic phase, including interventricular septum (IVSD), left ventricular posterior wall thickness (PWTED), left ventricular end diastolic dimension (LVEDD), and left ventricular end systolic dimension (LVESD). Three to five values for each measurement were acquired and averaged for evaluation. The LVEDD and LVESD were used to derive fractional shortening (FS) to measure left ventricular performance by the following equation: FS (%) ¼ [(LVEDD - LVESD)/LVEDD] x 100.

###### SPECT imaging/Annexin V Labeling

Recombinant human HYNIC annexin V prepared in *Escherichia coli* was obtained from the National Cancer Institute preclinical reagent program.11 To radiolabel the HYNIC-hydrazinonicotinamide-annexin V conjugate with 99mTc, 0.3 mL of saline containing approximately 1,000 MBq of 99mTc pertechnetate was added. Subsequently, 0.2 mL of freshly prepared stannous-tricine was added to the solution. The reaction vial was incubated for 15 to 20 minutes at room temperature. Radiochemical purity was determined chromatographically with instant thin-layer chromatography paper (Gelman Sciences, Ann Arbor, MI) using an acid citrate dextrose buffer (Sigma-Aldrich, St Louis, MO) as the mobile phase. Labeling efficiency consistently exceeded 92%, providing a specific radioactivity of 7.4 MBq/μg of protein.

In vivo SPECT was performed to quantify cell death in the heart. Rats were injected with 7 to 8 mCi 99mTc-HYNIC-annexin V and imaged using a FLEX XSPECT system (Gamma Medica-Ideas, Northridge, CA) 1 hour post injection. Two imaging methods were used: high-resolution dual-head single-pinhole imaging with 90 views and 30 s/view (total scanning time 45 minutes) or high detection efficiency five-pinhole imaging, with 90 views and 30 s/view. The data were reconstructed with an OS-EM-based three-dimensional pinhole reconstruction algorithm. CT acquisition (75 kVp, 0.24 mA, 512 projections, 0.1 s/projection) was also performed sequentially after SPECT imaging, and data were then registered with the corresponding SPECT images for anatomic delineation. **(Reference: Gabrielson 2008 Molecular Imaging)**

## Results and Discussion

### A: Drug Design

Most chemotherapeutics, currently used for TNBC treatments are non-specific or not targeted to any tumor specific biomarker^23^. Despite high off-target toxicity of chemotherapy, it is still a major player in the treatment of cancer. Particularly, true for triple negative breast cancer (TNBC) patients for whom no targeted therapy is not approved so far despite some of them are in clinical trials. The conventional targeted drugs are designed with a specific receptor in mind which is overexpressed in cancer cells compared to normal cells. For example, Cathepsin B is a well-studied tumor specific biomarker in various cancers including, breast, prostate, lung, liver, kidney brain and TNBC^10^. The prodrug approach of biological benign activity in circulation while, releasing the drug near target sites is an important part of the targeted drug design. Hence, both targeting and cleavage of the biologically active moieties are essential for high efficacy and safety of drugs. The conventional α-tocopheryl succinate (α-TOS) is a classic example of working very well *in vitro* and in certain immunocompromised *in vivo* tumor models while, failed in clinically relevant immunocompetent tumor models. For example, studies by Ireland et.al;^28^ showed that α-TOS was not only ineffective at the published doses, but resulted in severe side effects (e.g. loss of CD^3+^ T cells, organ fusing under peritoneum similar to standard toxin Terpenen-4-ol). Even high doses (471, 235 and 117 mg/Kg) did not achieve tumor regression *in vivo* (Fig 1a and b in ref 28). Although α-TOS has not been well studied in cardiomyocytes, α-TOS enhances cellular ROS levels in mitochondria which may help cancer cells to die while, it also induces cell death in cardiomyocytes similar to the mechanism for doxorubicin^29^. α-TOS is also sparingly soluble in water making it a difficult formulation for clinical purposes.

**Fig 1:**
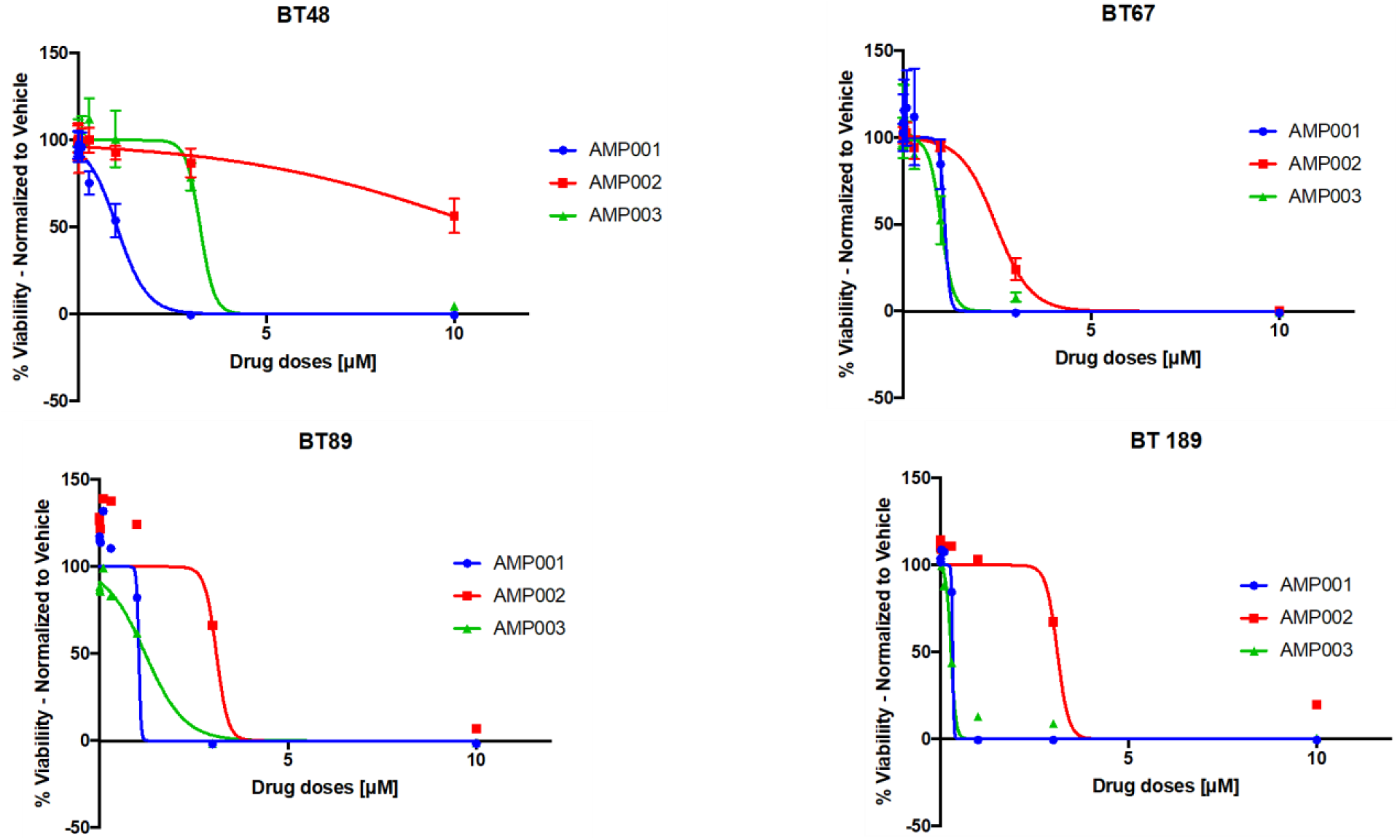
IC-50 values for AMP-001,002 & 003 in brain tumor stem cells determined using Cell-Titer-Glo Luminescent Cell viability assay, n= 4, p <0.002.

The lack of clinical product until now, for α-TOS, has provided impetus for us to redesign drugs to include a) <u>targeting/cleavable linker</u> to release the drug near tumor which is expected to reduce the off-target toxicity, b) <u>pegylation</u> to make it water soluble/bioavailable at higher doses and to keep it intact as a pro-drug in blood circulation for a longer time so that permeable angiogenic sprouts can uptake the drug^30^. Valine-Citrulline is a dipeptide linker which is known to be cleaved by Cathepsin B. Hence, AMP-001 is designed to conjugate valine citrulline through pegylated units. AMP-002 and AMP-003 were designed with ethereal and ester link respectively without pegylation.

### B: Selectivity

Targeting is essential to avoid side effects of the drug which was successfully adopted by bringing tumor specific biomarker Cathepsin B to which AMP class of drugs bind and get cleaved at the citrulline bond (see arrow in Scheme 1). The selectivity of AMP compounds for cancer cells is due to the binding of AMPs to Cathepsin B (which is overexpressed in several cancers) which cleaves and releases a) pegylated α-TOS, b) ethereal oxy tocopheryl succinate and c) α-TOS. We re-designed α-TOS as a new drug molecule, now targeted to cancer cells through a VC linker, and soluble due to addition of a poly ethylene glycol (PEG) moieties (5mg/10mL solubility). PEGylation offers significant and distinct pharmacological advantages over the unmodified α-TOS, including improved drug solubility, reduced dosage frequency, reduced toxicity, an extended circulating life, increased drug stability, enhanced protection from proteolytic degradation, decreased immunogenicity and antigenicity, and minimal loss of biological activity^47–49^. Thus, structurally and functionally AMP-001/003 are not equivalent of α-TOS and hence, are considered as novel drugs. Scheme 2 describes the selectivity of AMP drugs.

### B: Biological Activity

Extensive biological properties and potential mechanism of AMP derivatives have been recently published by Pandurangi et. al^31^. Preliminary studies were concentrated on TNBC cells and cancer stem cells (CSCs). In continuation with it, in this study, AMPs are studied in glioblastoma (GBM) brain tumor stem cells (BTSCs) and tumor cells derived from patients to expand the application of AAAPT technology to other cancers. Cancer stem cells (CSCs) are believed to represent approximately 1% of the tumor as a distinct population and cause relapse and metastasis for greater than 40 % of breast cancer patients^11^. Although the name CSC is an operational definition, these cells typically express higher levels of genes associated with stemness and epithelial to mesenchymal transition (EMT), which are involved in chemoresistance^17^. Chemotherapy can induce cell death only in bulk cancer cells and not in either CSCs or in low-responsive resistant cells. Amongst many proposed mechanisms of chemoresistance and relapse of cancer, cancer stem cells (CSCs) and cancer resistant cells (CRCs) either innate or drug induced stemness are found to be major culprits for therapy failure^1^. CSCs induction of resistance^33^, metastasis^34^ is imminent and found to be responsible for the high recurrence and making tumor cells refractory to the treatments . Conventional high dose chemotherapy and radiation therapy may even induce stemness in non-stem cancer cells^27^. GBM BTSCs encompass heterogenous populations of multipotent, self-renewing, and tumorigenic cells, which have been proposed to be at the root of therapeutic resistance and recurrence. We have shown in our earlier studies that AMPs target CSCs through inhibiting NF-kB pathway^31^. For GBM we selected brain tumor stem cells BT48, BT67, BT89 and BT 189 which grow under as neutrospheres (spheroid cell clusters). It is hypothesized that BTSCs can invade distant tissues far away from the tumor mass in GBM. First, the IC-50 values for AMP-001/003 are in the range 0.2-12 μM which are comparable and in fact lower than many standard chemotherapeutics (Fig 1, see Table 1 of ref). The potency of AMP-001/003 is important for a potential clinical translation. Fig 2 shows BTSCs invasion into 0.4mg/mL type I collagen surrounding matrix. Invasion was measured by quantifying the surface area of the cells as they invade the matrix over 24 hrs using image analysis. Untreated BTSCs in DMSO depicted uncontrolled invasion of BTSCs into the collagen matrix in 24 hrs (Fig 1 A & D) in contrast to AMP-001 treatment (0.5 μM, Fig 1, C & F) and standard drug Statt 3 inhibitor (positive control, 0.25 μM, Fig 2 B & E) which inhibited the invasion of BTSCs. The kinetics of invasion inhibition is graphically represented in Fig 2 G. Time-lapse videos of BTSC invasions are described in supplementary materials. In summary, we demonstrated the potency of AMP-001 to inhibit invasiveness of BT-48 chemo- and radioresistant BTSCs with efficiency comparable to an established inhibitor of BT-48 invasiveness, Stattic (Stat3 inhibitor).

**Fig 2:**
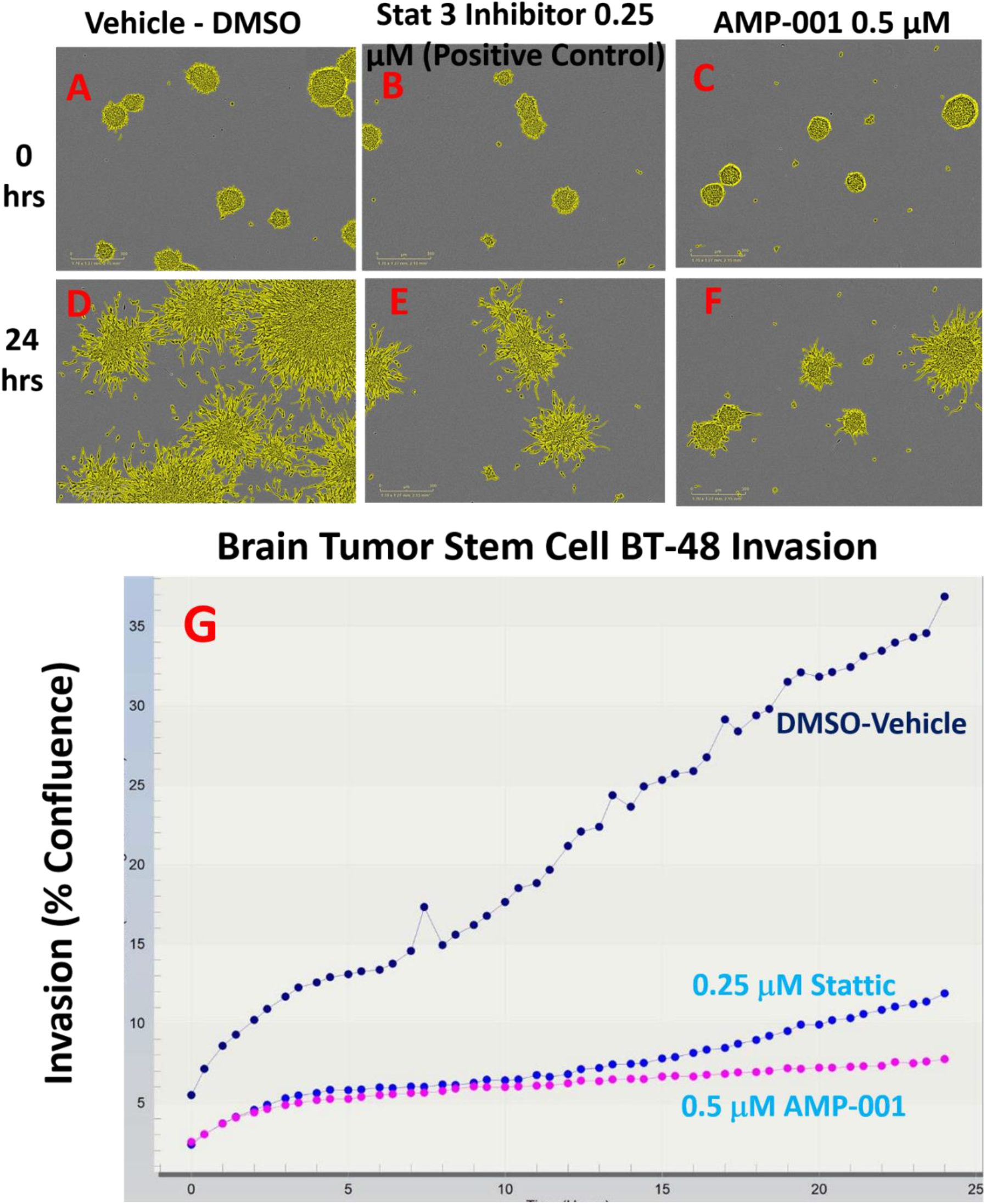
Live cell imaging of brain tumor stem cells (BTSC) neurosphere invasion into 0.4mg/mL type I collagen matrix the surrounding matrix with A) Vehicle Control, B) Drug Stattic (positive control) and C) AMP-001 at 0 and D-E-F at 24 hrs. n = 3, P < 0.003, Error bars represent standard error of mean (SEM), Scale bar 300 mm, G: Graphical representation.

### Synergy with Chemotherapy and Inhibitor Drugs

Chemotherapy is widely used for the treatment of most cancers^20^. However, low therapeutic index with a high dose related off-target toxicity limits their use in clinics. Since cancer is a group of diseases, recent studies have proposed targeting multiple targets requires a combined formulation with the other targeted drugs at adjuvant or neoadjuvant settings. Such an approach requires synergy between the two proposed combined formulation. In continuing with the synergistic potential of AAAPT, we further extended the studies to patient derived tumor cells which reflects the true heterogenous nature of tumors. Basal-like breast cancer is the most common subtype of TNBC prevalent, particularly in African Americans who had worst clinical outcomes in terms of relapse and survival. The characteristics of basal like breast cancer is the loss of PTEN (Phosphatase and Tensin homolog) in patient population which activates PI3K activation leading cancer cells to desensitize themselves to intervention. Alternate pathways to sensitize these cells to chemotherapy may benefit these subsets of breast cancer patients. Patient derived xenograft models of basal-like breast cancer cell lines (WU-BC3) were developed by Ma et.al^35^. which were treated with AMP-001 to see if the heterogeneity of cells has any effect on the synergistic potential of AMP-001. WU-BC3 cancer cells derived from patients was assessed for the loss of PTEN expression which is most common in many patients as described by Ma et.al^31^. Photomicrographs of WU-BC3 GFP-knockdown Human induced mice (Him3) triple negative breast cancer cells showed distinct morphology for most cancer cells when treated with low doses of AMP-001 alone or low doxorubicin (2 μM, Fig 3 A) which is another front-line treatment for TNBC. However, as the concentration of AMP-001 increases (12 and 25 μM) the combination of AMP-001 with Doxorubicin resulted in significant increase cancer cell death compared to individual drugs. Recent reports on the same WU-BC3 cells, PTEN knockdown led to a more dramatic reduction in cell proliferation and tumor growth inhibition in response to pan-AKT inhibitors both *in vitro* and *in vivo*. Similar results were obtained for WU-BC3 PTEN knockdown Human induced mice (Him3) triple negative breast cancer cells where photomicrographs showed synergistic cell death for the combination compared to individual drugs.

**Fig 3:**
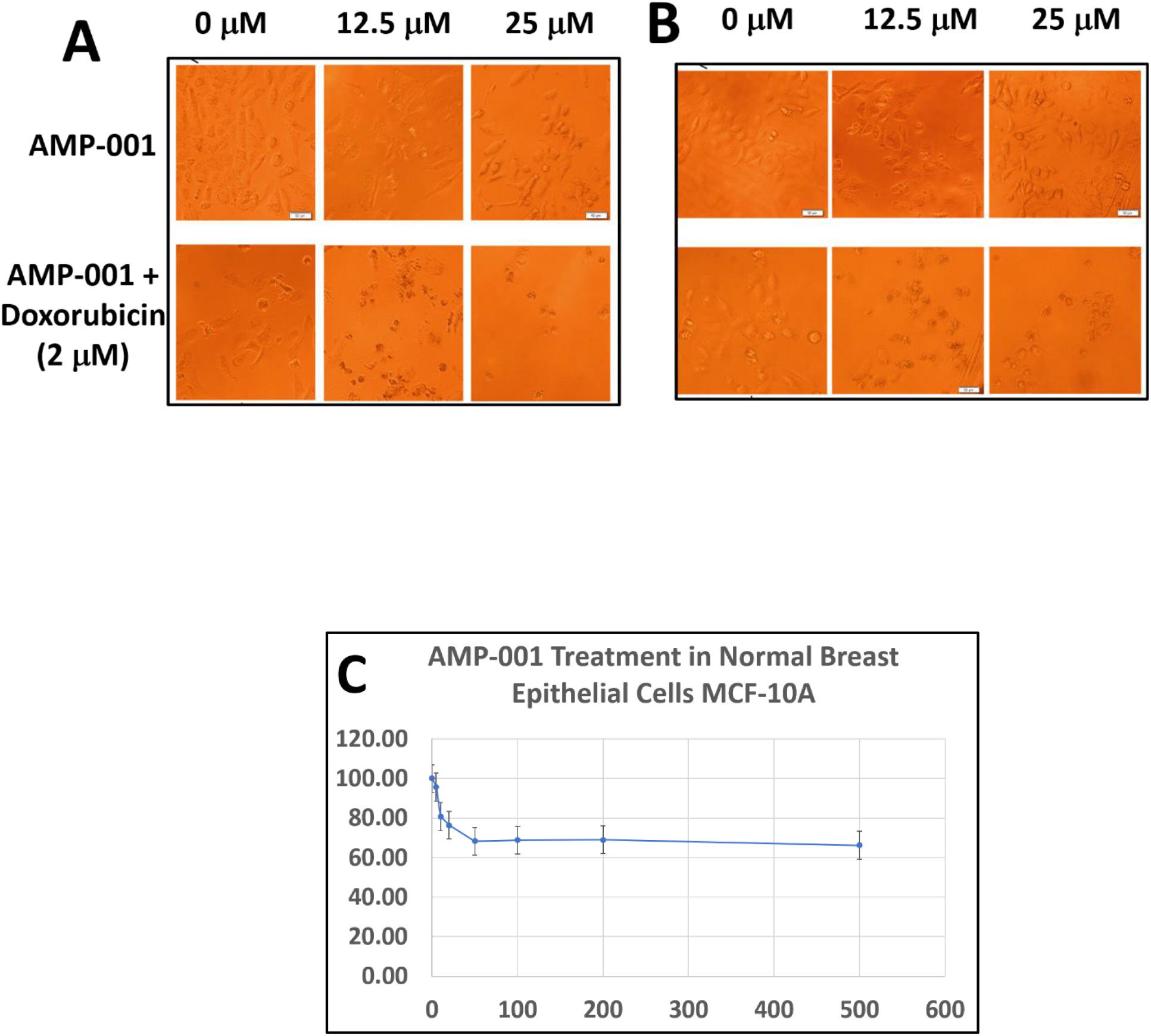
Synergistic apoptosis induction by AMP-001 in A) GFP-knockdown Human Induced Mice (Him3) triple negative breast cancer cells (WU-BC3), B) PTEN knocked down Him3, C) IC-50 of AMP-001 in normal breast epithelial cells MCF-10A, n =5, p < 0.003.

### Potency of AMP-001 in inducing tumor regression and enhancing the efficacy of doxorubicin at lower doses with no cardiotoxicity

We reported earlier the tumor regression by AMP-001 in TNBC mouse model without toxicity to off-target organs (e.g., heart, liver, and kidney)^31^. In order to extend the synergistic potential of AMP-001 from *in vitro* to *in vivo*, we developed rat TNBC model where we could monitor the toxicity of AMP-001 in non-target organs including heart which is mostly affected organ when treated with Doxorubicin clinically. Both noninvasive SPECT and Ultrasound imaging were used to assess the cardiotoxicity thorough cell death marker (SPECT) and ejection fraction respectively. The results revealed that Doxorubicin, (21mg/Kg) despite regressed tumor volume, it reduced the ejection fraction significantly from normal 65 % to 45 % (Fig 4 A & C). However, the combination of AMP-001 (150mg/Kg, i.p.) with 50 % reduced maximum tolerated dose (MTD) of Doxorubicin (10.5 mg/Kg, i.p.) achieved significant tumor regression (Fig 4 D, <u>*1850 mm*^*3*^ *to 280 mm*^*3*^</u>) with no cardiotoxicity compared to the MTD of Doxorubicin (i.e., 21mg/Kg, tumor regression, but cardiotoxic). Ultrasound imaging showed a ventricular dysfunction for Doxorubicin alone (EF < 45 %, Fig 4A) while AMP-001 with Doxorubicin combination retained ventricular function (EF ~65%, Fig 4, B-C).These data were corroborated with whole body SPECT imaging of cell death using 99mTc-Annexin which showed insignificant cell death in non-target organs for the combination (Fig 4, E1), while Doxorubicin alone showed significant cell death (Fig 4, E2).

**Fig 4:**
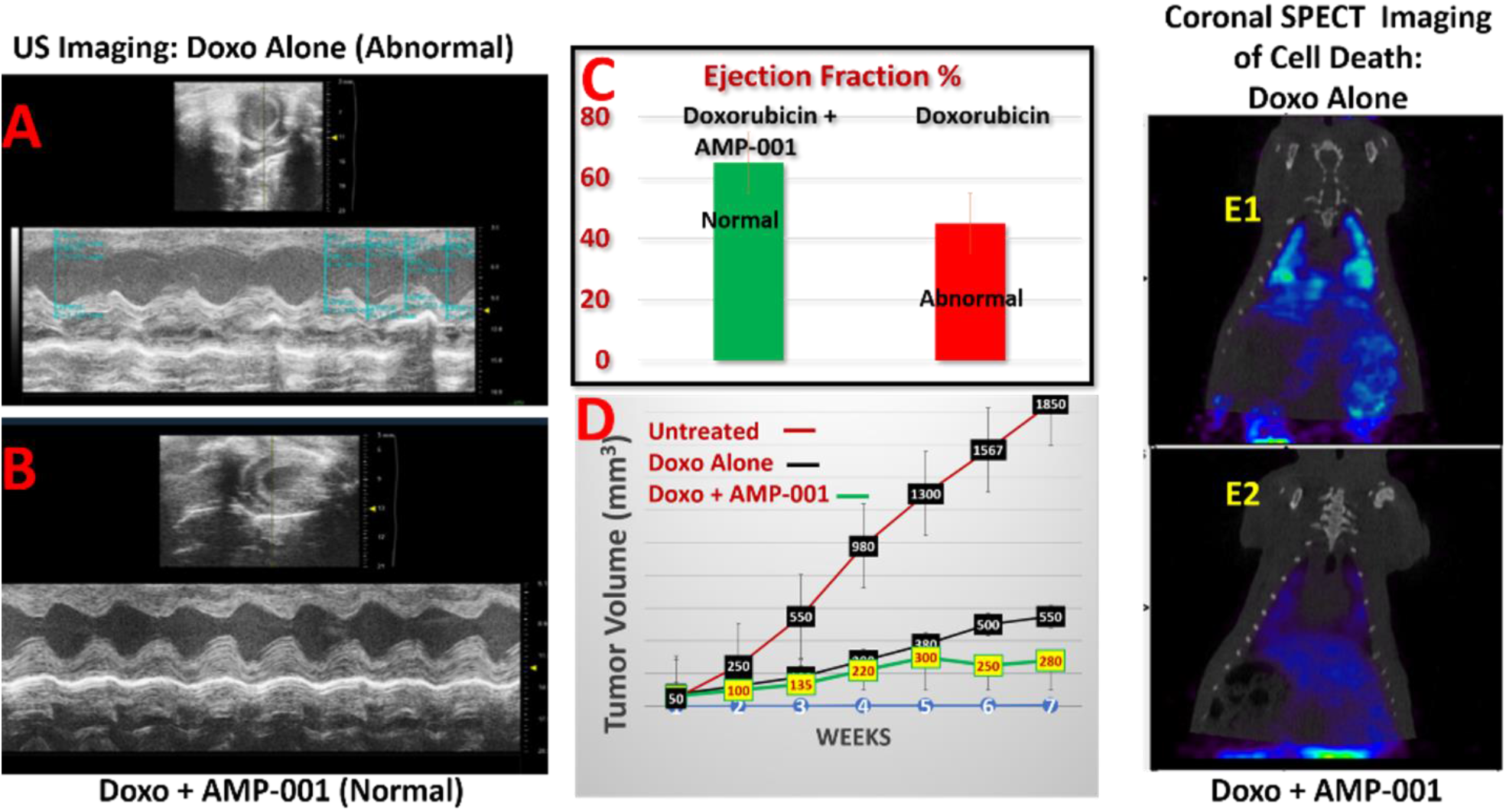
Effect of combination of AMP-001 and Doxorubicin on the tumor regression and cardiotoxicity in triple negative breast cancer rat model; A: Ultra sound imaging of the rat heart with a ventricular dysfunction, B: Reversal of ventricular dysfunction by the combination of AMP-001 and 50 % of Doxorubicin therapeutic dose, C; Graphical representation of ejection fraction for the combination of AMP-001 and 50% Doxorubicin (> 65%) compared to Doxorubicin alone, E: SPECT whole body imaging of cell death of TNBC rat using 99mTc-Annexin, E1: Doxorubicin alone Vs E2: Combination of AMP-001+ Doxorubicin.

The *in vivo* data were corroborated with data on pluripotent stem cell induced cardiomyocytes. Adult human induced pluripotent stem cell-derived (iPSC) cardiomyocyte technology offers the opportunity to accelerate the development of new therapeutic agents by providing a relevant human target for efficacy and safety that requires small amounts of test drug early in the research and development process. In other words, adult iPSC cardiomyocytes display normal cardiac physiology and electrophysiology, offering the potential to serve as a human-based cell model for safety pharmacology and cardiotoxicity screening. All iPSC cardiomyocytes were no longer attached to the cell plate and numerous cell fragments were suspended in the cell media for 10 μM doxorubicin while, AMP-001did not show the signs of toxicity even at 100 μM. After 24 hours exposure to AMP Conjugate all iPSCs were attached to the bottom of the cell well and contracting. These data demonstrate the IC-50 for AMP -001 for cardiomyocytes may be greater than 100 μM. Photomicrograph of normal, beating induced pluripotent stem cell cardiomyocytes (iPSCs) in 0.8% DMSO is shown in Fig 5A which is similar to the photomicrograph for AMP-001 at 200 mM (Fig 5B) conforming the low toxicity of AMP-001. In contrast, Doxorubicin showed morphological changes akin to apoptosis at 10 μM (Fig 5C). The IC-50 of doxorubicin for iPSCs was found to be 9.6 μM (Fig 5D). Concentration dependent viability assay for AMP-001 showed that iPSCs are viable even at 200 μM.

**Fig 5:**
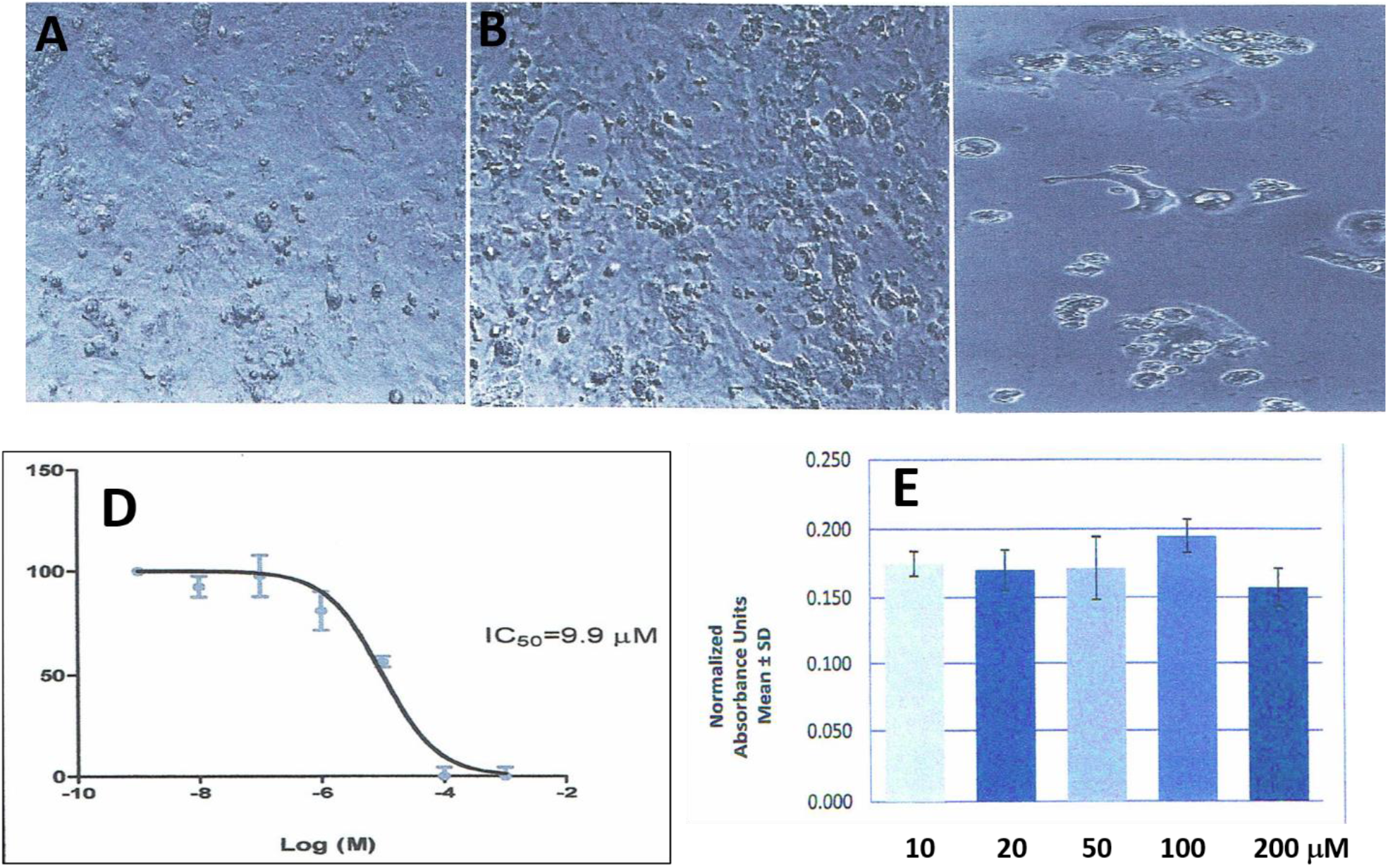
A: Photomicrograph of normal, beating induced pluripotent stem cell cardiomyocytes (iPSCs) in 0.8% DMSO, B: 200 μM of AMP-001, C: 10 μM Doxorubicin, D: IC-50 value for Doxorubicin in iPSCs, E: Concentration dependent viability for AMP-001 in iPSCs.

## Conclusions

Targeting drugs to specific biological sites requires the identification of a tumor specific biomarker which may help monitoring the efficacy from preclinical to clinical phase. AAAPT technology incorporates both targeting and cleavable linker technologies using tumor specific biomarker Cathepsin B and dipeptide valine citrulline linker (VC) which is cleaved by Cathepsin B. Using such drug designs, variety of drugs were synthesized with tocopheryl as a scaffold. Particularly, one unique AMP-001 conjugated Doxorubicin (AMP-004) is unique with two cleavable linkers hydrazone and VC releasing two warheads (AMP-001 and Doxorubicin) in vivo. All the leading drugs showed synergy with Doxorubicin which is the standard care of treatment, particularly for triple negative breast cancer treatment. AMP-001 also synergized with patient derived TNBC cells which are more heterogenous compared to cultured TNBC cells. Our studies indicated a potential for AMP-001 to use it as a neoadjuvant to Doxorubicin to increase the therapeutic index of Doxorubicin. The synergistic potential of AMP-001 with Doxorubicin enabled to reduce the therapeutic dose by 50 % which in turn reduced the cardiotoxicity associated with Doxorubicin. Further studies are required to extend the results to patient derived xenograft which mimics human tumor.

## Notes

### Competing Interest Statement

The authors have declared no competing interest.

